# Investigating the influence of iron parameters in regulating platelet recovery in Apheresis Platelet Donors

**DOI:** 10.1101/2024.08.22.609101

**Authors:** Ranita De, Leo Stephen, Vikram Mathews, Biju George, Eunice Sindhuvi Edison

## Abstract

**Introduction:** The present study investigates changes in haematological parameters in the donor during apheresis and following donation. Effect of iron parameters in influencing platelet recovery was studied.

**Methods:** Blood samples were collected in EDTA tubes for analyses of haematological parameters. Serum samples were isolated for estimation of iron parameters.

**Results:** Hb, RBC count and Hct increased during apheresis, but no significant changes were observed after 48 hrs post donation. Platelet recovery rates increased from 78 % after 48 hours to > 100% on day 10 (p < 0.001). Ferritin, serum iron & Transferrin saturation initially increased and subsequently declined from day 7 post donation, which may influence platelet recovery. Recovery rates were also higher in donors with lower iron stores.

**Conclusions:** Changes in haematological parameters during plateletpheresis are transient and optimal time required for platelet recovery is dependent on pre-donation counts. Recovery of platelet counts may be positively regulated by donor iron parameters.

## Introduction

The rising demand for platelet transfusions to treat patients affected with malignant and benign illnesses, those requiring organ/bone marrow transplants, surgical interventions or victims of traumatic injuries has made plateletpheresis one of the cornerstones of modern transfusion practice [1]. “Single Donor Platelets” (SDP) are preferred over Random Donor Platelets (RDP) as they have lower risk of infections and alloimmunization [2]. Different techniques have emerged since 1970’s to obtain Apheresis Platelet Concentrates (APC) from individual donors [3]. Although efficiency of platelet collection has increased in modern automated apheresis platforms, concerns regarding adverse reactions in plateletpheresis donors have also emerged [4].

To maintain donor platelet counts, the US Food and Drug Administration (FDA) had limited the number of donations to 12 per year, with a maximum of 2 donations per week and a minimum interval of 48hrs between successive donations. This was subsequently revised to permit up to 24 donations per year in 1988, and later restricted to donation of 24 components/year witha maximum limit of 3 components per apheresis procedure in 2005 [5]. Common haematological parameters have been reported to be adversely affected post plateletpheresis [6]. However, these changes may be transient, as no clinically relevant decrease in platelet counts has been observed in repeat platelet donors, when they donated within the limit set by the FDA. Some adverse effects on haematopoiesis have been observed in donors, undergoing multiple plateletpheresis procedures [8]. Thus, the effects of plateletpheresis on donor haematopoiesis remains contradictory and most of these studies have been carried out in the west.

Although recovery of platelet counts following apheresis has been studied, possible mechanisms responsible for the same are poorly elucidated. A close relationship has been documented between body iron stores and platelet counts, with thrombocytosis being a common complication in some patients affected by iron deficiency anemia [9]. Significance of iron deficiency in regulating thrombocytosis has been studied in animal models as well. For instance, R. Evstatiev et al. observed that young Sprague–Dawley rats fed on an iron deficient diet for a few weeks, showed increased platelet counts [10]. Some mechanistic insights into the role of iron deficiency in modulating megakaryopoiesis were obtained recently by Juliana Xavier-Ferrucio *et al.* They observed that mice in which the gene encoding the transmembrane serine protease 6 was knocked out (Tmprss6-/-) exhibited thrombocytosis, as high levels of hepcidin lead to systemic iron deficiency in these mice [11].

The present study investigates the changes in different haematological parameters in platelet donors during the apheresis procedure (within 30 minutes after initiation) as well as at 48 hours post donation. Optimal time required for recovery of platelet counts to baseline and potential significance of iron parameters in influencing the same was also studied.

## Materials and Methods

### 1. Settings and study design

This prospective observational study was conducted in the department of Haematology at a tertiary care hospital, in Vellore, Tamil Nadu in India, between May, 2021 and April, 2022. Consenting donors aged between 18-55 years were recruited and the study was approved by the Institutional Research and Ethics Committee. 32 healthy plateletpheresis donors eligible for donation, as per the criteria established by the Drug Controller General of India and the Director of General Health Services (DGHS) guidelines were enrolled in the study [12]. Donors were subjected to clinical examination and then those willing to donate by apheresis method were requested to complete the appropriate questionnaire form. They were also monitored for any adverse events during apheresis. Pre-donation samples from every donor were subjected to a complete blood count (CBC) using Yumizen H500 (HORIBA Medical) haematology analyser and screened for transfusion transmissible infections (TTI), as per departmental standard operating procedure.

### 2. Apheresis instrument & sample collection

The plateletpheresis procedure was performed with the Haemonetics MCS Plus cell separator using single-needle closed-system apheresis kits for SDP collection. Acidified citrate dextrose(ACD) was used as the anticoagulant in the ratio of 1:15 and maximum inlet rate varied from 60-80 ml/min. The target yield was set at 3.5 × 10^11^ platelets per unit. About 4 ml of peripheral blood was collected from donors in Ethylene Diamine Tetra Acetic Acid (EDTA) within 30 minutes after the start of the procedure and at 48 hours post donation (n=32). Additionally peripheral blood was also collected at 96 hours, day-7 and on day-10 post donation in some local voluntary donors who were willing to donate additional follow up samples (n=10). The latter samples were collected to investigate optimal time required for platelet recovery till baseline and effect of iron parameters in influencing the same.

#### 3. Estimation of haematological & iron parameters

Routine haematological and platelet related parameters were measured using Yumizen H500 (HORIBA Medical) haematology analyser. Serum samples isolated from peripheral blood of donors were stored at -80^0^ C for further analyses of iron parameters. Serum ferritin levels were determined on *Siemens ADVIA Centaur analyser* by two-site sandwich chemiluminescence immunoassay. Serum hepcidin levels were estimated by ELISA method (Hepcidin 25 bioactive ELISA, DRG International, Inc.). sTfR levels were determined on Roche *Cobas 8000* modular *analyser* using Tina-quant Roche kit by Immunoturbidimetric method. Serum iron levels and unsaturated iron-binding capacity (UIBC) were estimated by automated analysers.

Total iron-binding capacity (TIBC) was calculated using the formulae = UIBC + Serum iron Percentage of Transferrin saturation (TF%) was calculated by the formulae = Serum iron / TIBC ×100

### 4. Statistical analyses

Quantitative variables were reported using Mean ± Standard deviation (SD) or Median and interquartile range (IQR), according to the characteristics of the distribution of the data. For qualitative variables, number and percentages were used. Student’s t test was used to compare different haematological parameters between baseline levels and certain time points during apheresis and post donation. Comparison of platelet counts and iron parameters of donors at different time points was done by ANOVA (Graphpad Prism 8.0.1). Donors were divided into different groups, based on their iron content –

A. Ferritin low group (Ferritin-L) and ferritin high group (Ferritin-H) – donors with serum ferritin <group median and > group median, respectively.
B. sTfR low group (sTfR-L) and sTfR high group (sTfR-H) – donors with sTfR < group median and > group median, respectively.
C. Hepcidin low group (Hepcidin-L) and hepcidin high group (Hepcidin-H) – donors with serum hepcidin < group median and > group median, respectively.

Platelet recovery rates following apheresis were compared between donors of the three groups mentioned above.

Recovery of platelet count (in %) with respect to baseline count at different time points was calculated by the formula = Platelet count at specific time/ baseline platelet count × 100

For all tests, p-value of < 0.05 was considered as being statistically significant.

## 5. Results

### 5.1. Changes in haematological parameters during apheresis and post donation

The present study included 32 healthy male donors, who underwent plateletpheresis on the Haemonetics MCS Plus cell separator. Peripheral blood was collected before donation, during apheresis (within 30 minutes post initiation) and at 48 hours post donation. Their median age was 31 years (18-55). The mean Hb increased from baseline value of 14.4 ± 1.2 g/dL to 15.01 ± 1.1 g/dL within 30 minutes post initiation of the procedure. The mean RBC count increased from 5.03 ± 0.51 × 10^6^/μl to 5.27 ± 0.52 × 10^6^/μl within the same time. Similar trends were also observed in Hct which increased from 43.8 ± 2.9 % to 45.18 ± 3.6 % within the same time interval, post initiation of the apheresis procedure. However, these changes were not statistically significant and transient. After 48 hours post donation, no significant changes were observed in mean Hb (14.7 ± 0.93 g/dL), RBC count (5.21 ± 0.79 × 10^6^/μl) as well as mean Hct (45.03 ± 5.8 %) when compared to pre-donation levels.

Platelet counts significantly decreased from 2.65 ± 0.8 × 10^5^/μl to 2.02 ± 0.7 ×10^5^/μl within 30 minutes (p= 0.002) which further decreased to 1.99 ± 0.5 × 10^6^/μl at 48 hours post donation (p< 0.001). WBC counts also declined from 7.96 ± 2.1 ×10^3^/μl to 7.23 ± 1.8 ×10^3^/μl within 30 minutes post apheresis initiation (p= 0.143) and were measured as 7.8 ± 2.0 ×10^3^/μl at 48 hours post donation. No significant changes were observed in other platelet related parameters, MPV and PDW between baseline levels and during apheresis. However, MPV increased significantly from baseline value of 9.01 ± 1.06 (fl) to 9.6 ± 1.1 (fl) (p= 0.03) at 48 hours post donation. This corroborates with significantly decreased platelet counts observed at 48 hours post donation,as an inverse correlation between these parameters has been reported in individuals with low iron stores [13]. These changes in different haematological parameters have been summarized in Table 1.

**Table 1:** Summary of changes in haematological parameters during apheresis and post platelet donation (n = 32)

| Parameter | Baseline | During<br>apheresis | 48 hrs post<br>donation | Baseline & during<br>apheresis<br>*p*-value | Baseline & 48 hrs<br>post donation<br>*p*-value |
| --- | --- | --- | --- | --- | --- |
| **RBC (×10<sup>6</sup>/μl)** | 5.03 ± 0.51 | 5.27 ± 0.52 | 5.21 ± 0.7 | 0.075 | 0.297 |
| **Hb (g/dL)** | 14.4 ± 1.2 | 15.01 ± 1.1 | 14.7 ± 0.9 | 0.072 | 0.389 |
| **Hct (%)** | 43.8 ± 2.9 | 45.1 ± 3.6 | 45.0 ± 5.8 | 0.112 | 0.310 |
| **MCV (fl)** | 86.3 ± 6.1 | 86.9 ± 5.8 | 86.7 ± 5.7 | 0.703 | 0.774 |
| **WBC (×10<sup>3</sup>/μl)** | 7.96 ± 2.1 | 7.2 ± 1.8 | 7.8 ± 2.0 | 0.143 | 0.754 |
| **Plt count (×10<sup>5</sup>/μl)** | 2.65 ± 0.8 | 2.02 ± 0.7 | 1.99 ± 0.5 | 0.002 | < 0.001 |
| **PDW (fl)** | 12.9 ± 2.6 | 13.5 ± 2.8 | 13.9 ± 3.0 | 0.418 | 0.156 |
| **MPV (fl)** | 9.01 ± 1.06 | 9.5 ± 1.1 | 9.6 ± 1.1 | 0.060 | 0.033 |

\*Mean ± SD are indicated for different parameters

### 5.2. Recovery of platelet counts at different time points following apheresis

The recovery of platelet counts post donation was studied in some donors, who volunteered to give additional samples at subsequent time points (n=10). Platelet counts progressively increased following an initial decline. The mean baseline platelet count of 2.64 ± 0.5 ×10^5^/μl decreased significantly to 2.02 ± 0.4 ×10^5^/μl at 48 hours (p= 0.01), following which it increased to 2.22 ± 0.43 ×10^5^/μl at 96 hours post donation. Further increase in mean platelet counts were observed on day-7 (2.53 ± 0.54 ×10^5^/μl) and on day-10 (2.81 ± 0.61 ×10^5^/μl). Thus, the donor platelet counts had recovered by 78 % by 48 hours, when compared to the baseline. By day-7, the mean platelet counts increased by 19.3% compared to 48 hrs (p = 0.03), which further increased significantly beyond baseline counts by day-10 (p < 0.001). In fact, an excess of 8 % above the baseline count was measured on day-10 post donation. The platelet recovery rates (in %), with respect to baseline have been depicted in Figure 1.

**Figure 1:**
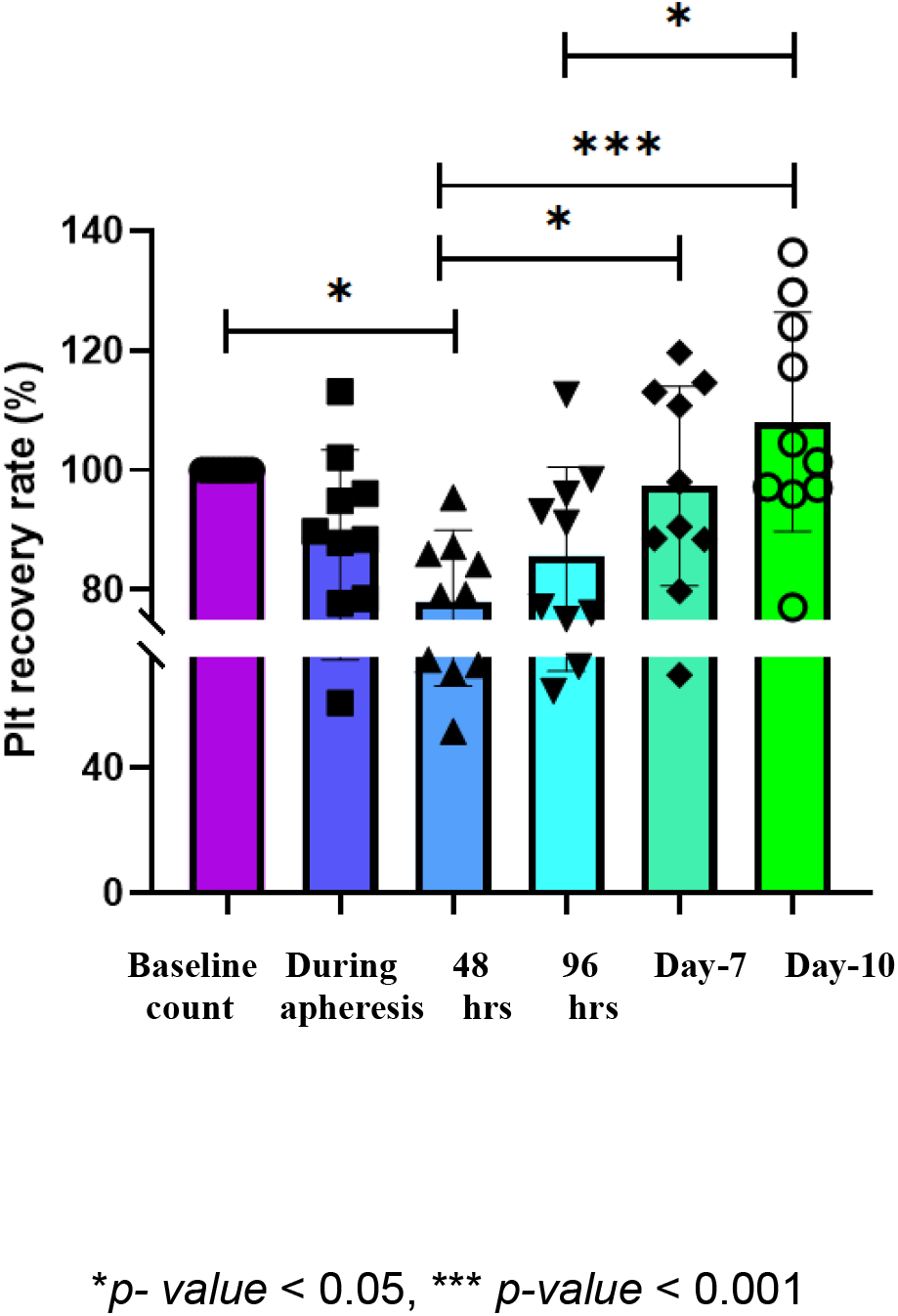
Comparison of platelet recovery rates in donors at different time points following apheresis (n=10) Units of platelet recovery rate, (compared with baseline) - % \**p-value* < 0.05, *** *p-value* < 0.001

#### 5.3. Significance of iron parameters in influencing platelet recovery

Serum ferritin levels increased from 99.8 ng/ml (26.7-270.4 ng/ml) within 30 minutes post initiation of apheresis to 112 ng/ml (41.8-307.6 ng/ml) at 96 hours post donation. This may be due to decrease in median serum hepcidin levels from 9.6 ng/ml (0.9-17.3 ng/ml) at 30 minutes post initiation to 6.36 ng/ml (0.6-18 ng/ml) at 96 hours following donation. Hepcidin levels subsequently increased to 10.16 ng/ml (0.82-17.5 ng/ml) on day 10. This may be responsible for decline in serum ferritin levels to 82.9 ng/ml (17.5-280.3 ng/ml) on day 10 post donation. These changes were not significant.

Similar trends were observed when serum iron levels were measured in these donors. Median serum iron level increased from 85 µg/dL (57-116 µg/dL) at 30 minutes post initiation to 92 µg/dL (43-128 µg/dL) at 96 hours post donation. On day 10 post donation the median serum iron levels declined to 85.5 µg/dL (58-113 µg/dL). As iron stores increased initially, platelet recovery rates decreased to 78 % at 48 hours post donation, compared to baseline platelet counts. They subsequently increased to 85.6% after 96 hours and recovered to > 100% on day 10 post donation, while serum ferritin and iron declined after day 7.

Similarly, serum TF% increased from 23.7± 6.4 % at 30 minutes post initiation at subsequent time points, eventually decreasing to 21.8 ± 5.02 % on day-10. sTfR levels declined from 3.24 ± 1.0 mg/l during apheresis to 3.06 ± 0.8 mg/l at 96 hours post donation. It gradually increased to 3.16 ± 0.9 mg/l on day 10, correlating with decreasing iron stores. These changes were not statistically significant. Variation in iron parameters in these donors at different time points post apheresis is summarized in Table 2.

**Table 2:** Comparison of changes in iron parameters at different time points following apheresis in platelet donors (n = 10)

| Iron parameter | During apheresis | 48 hrs post donation | 96 hrs post donation | Day-7 post donation | Day-10 post donation |
| --- | --- | --- | --- | --- | --- |
| **Serum Ferritin**<br>(ng/ml) | 99.8<br>(26.7-270.4) | 100<br>(16.4-321.5) | 112<br>(39.9-307.6) | 88.7<br>(15.1-294.5) | 82.8<br>(17.5-283) |
| **Serum Iron**<br>(µg/dL) | 85<br>(57-116) | 82<br>(62-182) | 92<br>(43-128) | 88.5<br>(60-138) | 85.5<br>(58-113) |
| **Transferrin saturation (%)** | 23.78 ± 6.4 | 25.95 ± 8.7 | 26.26 ± 10.4 | 22.37 ± 8.3 | 21.81 ± 5.02 |
| **Hepcidin**<br>(ng/ml) | 9.61<br>(0.9-17.3) | 6.48<br>(0.4-14.6) | 6.36<br>(0.6-18) | 7.34<br>(0.67-14.6) | 10.16<br>(0.82-17.5) |
| **sTfR**<br>(mg/L) | 3.24 ± 1.05 | 3.05 ± 0.86 | 3.06 ± 0.89 | 3.09 ± 0.92 | 3.15 ± 0.98 |

\*Mean ± SD and Median (IQR) for different iron parameters are indicated.

When platelet recovery rates were studied in Ferritin-L and Ferritin-H groups i.e. donors with ferritin < median (99.8 ng/ml) and > median respectively, platelet counts recovered to > 100 % of baseline by day 10 in both groups. However, donors in the Ferritin-L group showed greater platelet recovery on day 7 and 10, as compared to donors in the Ferritin-H group. Platelet recovery rates were also higher in donors with hepcidin > group median (9.61 ng/ml) i.e. (Hepcidin-H) vs donors having hepcidin levels < median value (Hepcidin-L). Similar rates of platelet recovery were observed between donors with sTfR < group median (3.47 mg/l) i.e. (sTfR-L) and those having sTfR levels > median value (sTfR-H). Changes in platelet recovery rates in donors categorized into different groups based on iron parameters have been presented in Figure 2.

**Figure 2:**
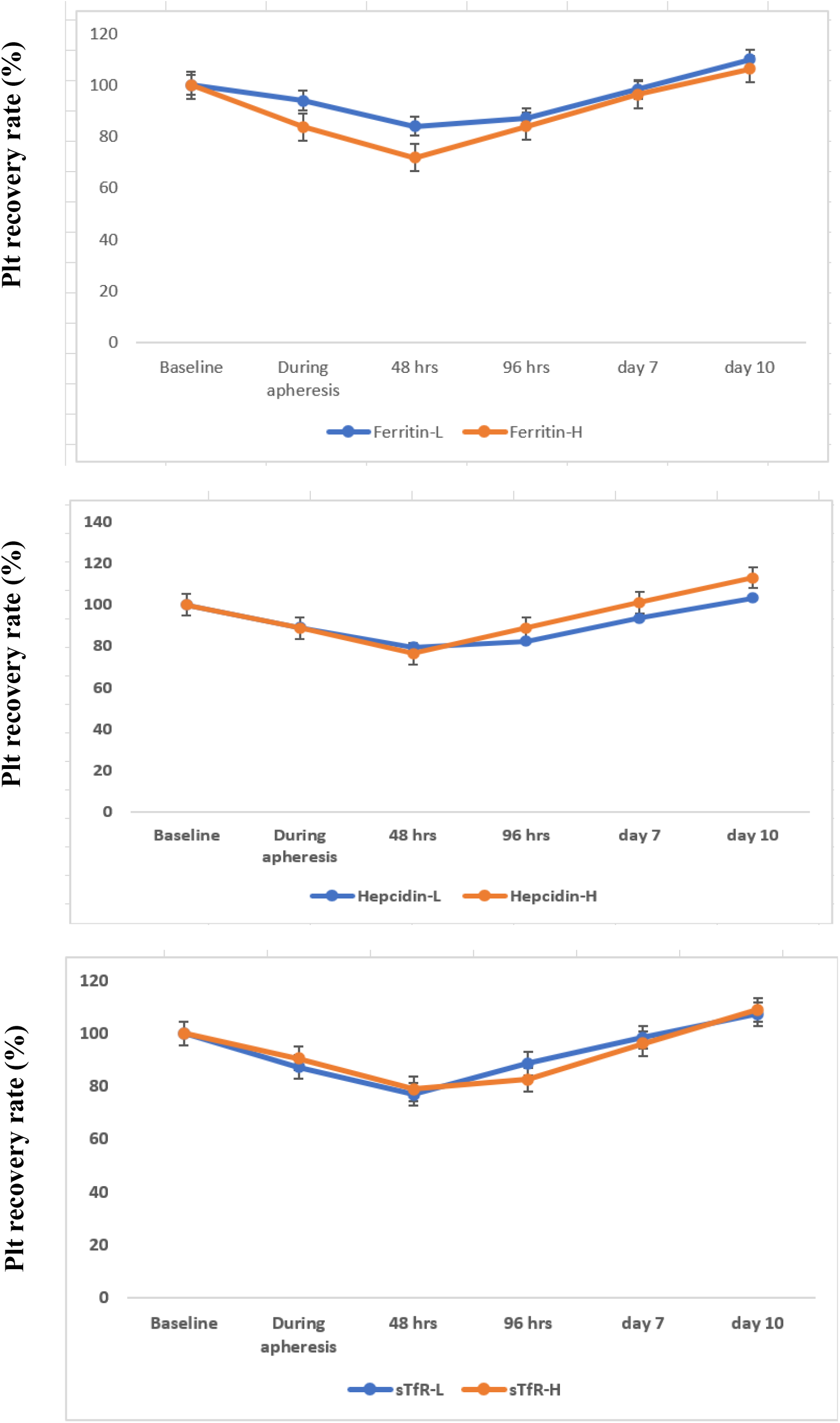
Comparison of platelet recovery rates in donors categorized into different groups based on iron parameters (n=10) Units of platelet recovery rate, (compared with baseline) - % Abbreviations: Ferritin-L and Ferritin-H – donors with serum ferritin < group median (99.8 ng/ml) and > group median, respectively; sTfR-L and sTfR-H – donors with sTfR < group median (3.47 mg/l) and > group median, respectively; Hepcidin-L and hepcidin-H – donors with serum hepcidin < group median (9.61 ng/ml) and > group median, respectively.

## 6. Discussion

The mean pre-donation platelet count observed in the present study (2.64 ± 0.5 ×10^5^/μl) conducted at CMC, Vellore in Tamil Nadu was similar to another study conducted in southern India by Suresh et al. (2.80 ± 0.55 ×10^5^/μl) [14]. Recent studies by Syal et al. and Rajput et al. from north India reported mean pre-donation platelet counts of 2.67 ± 0.5 ×10^5^/μl [15] and 2.39 ± 0.56 ×10^5^/μl [16] respectively. These studies investigated the effect of plateletpheresis on haematological parameters at different time points after donation.

Syal et al. reported a significant increase in Hb, Hct and RBC count when measured at 30 minutes post donation, when plateletpheresis was carried out on the Haemonetics MCS+ cell separator [15]. Similar increase in Hb and Hct were also reported by Sachdeva et al., when these parameters were measured post platelet donation [17]. In the present study, we observed that although an increase in Hb, RBC count and Hct were observed during apheresis i.e. within 30 minutes post initiation, these changes were not statistically significant and transient. As most past studies have looked at changes in haematological parameters immediately post apheresis, we wanted to investigate what happens to these parameters during apheresis which may lead to clinical manifestations post donation. However, at 48 hours after donation,there were no significant changes in Hb, RBC count and Hct compared to baseline levels. Suresh et al. reported significant decrease in Hb, Hct at 30 minutes post donation, when the Fresenius Kabi COM.TEC apheresis machine was used [14]. These findings were similar to an older study by Das et al. [6]. Recently Rajput et al. also reported non-significant decline in Hb, Hct and RBC count post donation, when apheresis was performed with the Amicus cell separation platform [16]. Infusion of anticoagulants and normal saline or mechanical hemolysis during the procedure may account for decrease in haematological parameters [6]. Different machines used for apheresis may also be responsible, as the void volume of the Haemonetics MCS+ is very less compared to other machines. Hence blood loss in the void volume is minimal at the end of apheresis, when performed with the former. This may explain increase in haematological parameters observed post donation by Syal et al. [15], as compared to other studies which reported decline in these parameters when alternate apheresis platforms were used [6],[16]. Mean WBC count decreased in our study from baseline count of 7.96 ± 2.1 ×10^3^/μl to 7.23 ± 1.8 ×10^3^/μl at 30 minutes post initiation of apheresis, this change was not significant.Earlier studies by Das et al. [6] and Suresh et al. [14] as well as a recent study by Syal et al. [15] reported significant decrease in WBC count, post donation. This may be explained by the fact that blood components collected by all automated cytapheresis contain donor leukocytes [15].

Decrease in donor platelet count post apheresis has many reported in many studies. Suresh et al. demonstrated that donors with baseline platelet count of 2.80 ± 0.55 lost 1.05 lakhs/µl of platelets within 30 minutes post donation [14]. Recent studies by Syal et al. and Rajput et al.reported similar platelet losses of 0.74 lakhs/ µl [15] and 0.72 lakhs/ µl [16] respectively, with target platelet yield of ≥ 3 × 10^11^ within the same time interval post donation. In the present study, donors with mean baseline count of 2.64 ± 0.55 lost 0.33 lakhs/ µl during apheresis i.e. within 30 minutes of its initiation. After 48 hours post donation, donors had lost 0.62 lakhs/ µl of platelets which declined to 0.11 lakhs/ µl on day-7 and recovered in excess of baseline count on day 14. Recovery of platelet counts till baseline studied by different researchers in the past have yielded contrasting results. Thokala et al. observed that platelet counts recovered to > 100% compared to baseline, on day-7 post donation in donors with mean platelet count between 1.5-2.75 ×10^5^/μl (n= 35). Where as in donors with higher mean platelet count of > 2.75-3.5 ×10^5^/μl (n=12) platelet counts recovered till baseline after 2 weeks following donation [18]. Similar results were observed in the present study, where in donors with relatively high mean platelet count of 2.64 ± 0.55 ×10^5^/μl showed recovery of > 95% of platelet counts by day 7, which increased to > 100% by day 10 post donation (n=10). A recent study by Rajput et. al. where 30 donors were followed up following plateletpheresis, indicated that platelet counts recovered to 94.2% of the pre-donation count (2.2 ± 0.4) within 48 hours [16]. Thus, the time required for recovery of platelet counts following donation seems to be dependent on initial pre-donation counts.

Possible mechanisms responsible for platelet recovery following apheresis were investigated in an older study by Wagner et al. They reported that serum thrombopoietin levels increased by day 1 following apheresis, in 12 healthy donors and remained elevated on day 7. The number of peripheral blood colony forming unit-megakaryocytes (CFU-Mk) also increased by day 4 post platelet donation. Thus, the recovery trend observed in platelet counts following apheresis may be due to changes in platelet progenitor cells, through alterations in serum TPO levels [19]. A close relationship has been documented between body iron stores and platelet counts, with thrombocytosis being a common complication of some patients affected with iron deficiency anemia [20]. In the present study, we investigated potential changes in iron parameters in apheresis donors at different time points post donation. Serum ferritin,serum iron as well as transferrin saturation levels gradually increased when measured within 30 minutes post initiation of apheresis, till 96 hours post donation. This may be facilitated by decrease in serum hepcidin levels during the same time interval. Platelet recovery rates on the other hand decreased from 88.8 % at 30 minutes post apheresis initiation to 85.6 % at 96 hours post donation. Following this, the recovery rate increased to >95 % on day 7 and reached > 100% on 10^th^ day after apheresis. Interestingly, serum ferritin, iron and transferrin saturation started reducing by day 7 and decreased further on day 10. This may be explained by increase in serum hepcidin levels by day 10. Thus, the linear trend of gradual increase in platelet counts observed after 7 days post apheresis, may be influenced by declining iron stores in these donors. These observations were also supported by greater platelet recovery rates in donors with ferritin content lesser than group median (99.8 ng/ml) vs donors having ferritin content > median value. Similarly, donors with hepcidin content > group median (9.61 ng/ml)possessing lower intracellular iron stores showed greater platelet recovery rates on day 7 as well as 10, compared to donors with hepcidin content < median value. The present study is the first one, to the best of our knowledge which investigates the potential significance of iron parameters in affecting recovery of platelet counts in plateletpheresis donors.

## 7. Conclusions

The variation in haematological parameters during apheresis is transient and no significant changes were observed in plateletpheresis donors at 48 hours after donation. Although platelet counts dropped significantly after 48 hours post donation, they recovered till base line counts by day 10. The time duration required for platelet recovery has yielded contrasting results in recent studies and appears shorter for donors with lower pre-donation counts. We observed that different iron parameters such as serum iron, ferritin and transferrin saturation initially increased and subsequently declined from day 7 post donation. This may influence the increasing rate of platelet recovery from > 95 % on day 7 to >100% of the baseline count by day 10. When donors were categorized by their median ferritin and hepcidin values, greater platelet recovery rates were also observed in donors having relatively lower ferritin and higher hepcidin (facilitating lower intracellular iron stores) than their counterparts. These observations indicate platelet recovery post apheresis may be positively regulated by donor iron levels. This may be investigated in a larger cohort to ascertain the significance of donor iron stores, in influencing recovery of platelet counts following plateletpheresis.

## Acknowledgements

RD performed experiments and analyses, wrote, and edited the manuscript. LS helped in sample collection and processing. VM and BG edited and reviewed the manuscript. ES designed the research proposal, analyzed the data, and reviewed the manuscript. All authors read and approved the final version of the manuscript.

## Data Availability Statement

Data will be available upon request, by the corresponding author.

